# 30×30 biodiversity gains rely on national coordination

**DOI:** 10.1101/2022.12.16.520787

**Authors:** Isaac Eckert, Andrea Brown, Dominique Caron, Federico Riva, Laura J. Pollock

**Affiliations:** Dept. of Biology, McGill University, 1205 Docteur Penfield Montréal, H3A 1B1, Québec, Canada; Québec Centre for Biodiversity Science, Montréal, Québec, Canada; Institute of Geography and Sustainability, University of Lausanne, Lausanne, Switzerland

## Abstract

Protecting 30% of land by 2030 is an invaluable opportunity to combat the ongoing biodiversity crisis, but critical questions remain regarding what biodiversity to prioritize, how to coordinate protection, and how to incorporate global change. Here, we evaluate how well different 30×30 expansion scenarios capture the climatically viable ranges of Canadian terrestrial vertebrates, plants, and butterflies. We find that national coordination protects vastly more biodiversity (65% of species; 40% of species-at-risk) than regional approaches, which safeguard at least 33% fewer species and 75% fewer species-at-risk. Whereas prioritizing different taxa or biodiversity facets (e.g., phylogenetic diversity) incur smaller trade-offs. Surprisingly, national priorities closely match transnational ones, indicating that national coordination could efficiently contribute to global targets while protecting Canada’s biodiversity at large.

## Main Text

Protected areas are pivotal to biodiversity conservation and increasing the amount of protected land is at the forefront of global conservation policy (*1*). Not all land however, is equal, and the success of international goals like protecting 30% of land by 2030 (30×30) depends both on our capacity to protect more land and our ability to capture essential biodiversity when we do (*2*). So far, protected areas have largely failed at this latter goal, due to historical principles (*3*), the uneven distribution of biodiversity (*4*), and the omission of biodiversity in spatial planning (*5*). Fortunately, decades of convergent scientific research, public support, and international momentum have culminated with ambitious area-based targets in the post-2020 Global Biodiversity Framework (GBF) and the commitment of over 100 countries to 30×30 (*1*), giving humanity a chance to prevent future biodiversity loss and potential erosion of critical ecosystem services. But to facilitate positive outcomes for nature, we first need to take stock of what is already protected, estimate what could be protected if we reach 30×30, and understand how different conservation priorities influence our ability to protect biodiversity into the future (*5*).

Prioritizing land for protection is a long-standing challenge in conservation, with solutions varying widely depending on conservation values and practical constraints of capital, tool, and data availability (*6*). Historically, protected areas were often founded to protect landscapes, not biodiversity (*3*). Where biodiversity has been considered, the focus has typically been on specific taxa (i.e., migratory birds) or species at-risk (*7*). Even recent approaches focused more on entire ecosystems, such as Key Biodiversity Areas, consider individual target species deemed important to global biodiversity (*8*). Such a focus can be useful, and understanding how conservation priorities differ across groups can inform strategy and uncover patterns (*9*). However, conservation assessments are often biased, incomplete, or lacking consideration of future factors like climate change (*10*) and the trade-offs for non-target taxa are often unquantified and unknown.

In contrast to focusing on specific taxa, spatial planning can now consider a broad range of biodiversity, due to advances in biodiversity science and the shrinking Wallacean shortfall (lack of knowledge of all species ranges) (*11*). Biodiversity indicators can now be mapped and evaluated for thousands of species (*12*) and functional and phylogenetic facets directly link species to functioning ecosystems, future option values, and millions of years of evolutionary change (*13*). With the proliferation of new indicators comes an even greater need to determine the inherent trade-offs associated with choosing one element of biodiversity to prioritize over another, and recent research has begun to explore the trade-offs between biodiversity and other facets (*13*). Further, the added urgency of climate change means that it is increasingly important to directly consider future climates in spatial planning (*15*). Early papers established a framework for doing this by prioritizing “win-win” areas within a species current range that remain climatically-viable into the future (*16*), but few studies have incorporated climatic viability alongside multiple conservation priorities.

Finally, even if biodiversity values and metrics are agreed upon, the scale at which protection is coordinated can influence spatial priorities and biodiversity outcomes. At the extreme, a global target requiring full transnational coordination would optimally protect biodiversity in the broadest sense (*13*). More realistic, however, is coordination at the national scale to reach national and transnational targets. But national coordination might not capture transnational priorities, and even if they do, national priorities could easily contrast with the often (nationally) uncoordinated, but locally effective, reality of regional conservation initiatives (*17*). For example, if jurisdictions focus exclusively on regionally important biodiversity, this could limit their contributions to national or global targets. Despite this, the concept of spatial representation (i.e., parochialism), where protected land is distributed evenly across political or ecological regions, was a substantial requirement in the pre-2020 Aichi Biodiversity Targets (*6*) and in the post-2020 GBF (*1*). Since conservation regularly happens at regional scales, spatial representation is already occurring to some extent (*18*). Surprisingly, the biodiversity trade-offs associated with transnational versus national versus regional strategies remain largely unknown, despite their central importance to targets like 30×30.

Here, we bring together recent advances in biodiversity modeling, spatial prioritization, and indicator development to assess how different approaches to reaching 30×30 incur trade-offs and impact our ability to protect biodiversity. Focusing on Canada, a country with considerable conservation potential (*19*), a rapidly changing climate (*20*), and a poor international conservation record regarding area-based targets (*21*), we modeled the distribution of terrestrial vertebrates (460 birds, 147 mammals, 90 amphibians & reptiles), plants (3378), and butterflies (190). Using these models, we evaluated Canada’s existing network of protected areas and identified land that would need to be protected to reach 30×30 while maximizing positive outcomes for biodiversity. We incorporated future climates by valuing “win-win” areas expected to be stable under climate change based on current and future projections of species distributions (under RCP8.5), down weighting future distributions to account for uncertainty (Fig.1) (*16*).

**Fig. 1.**
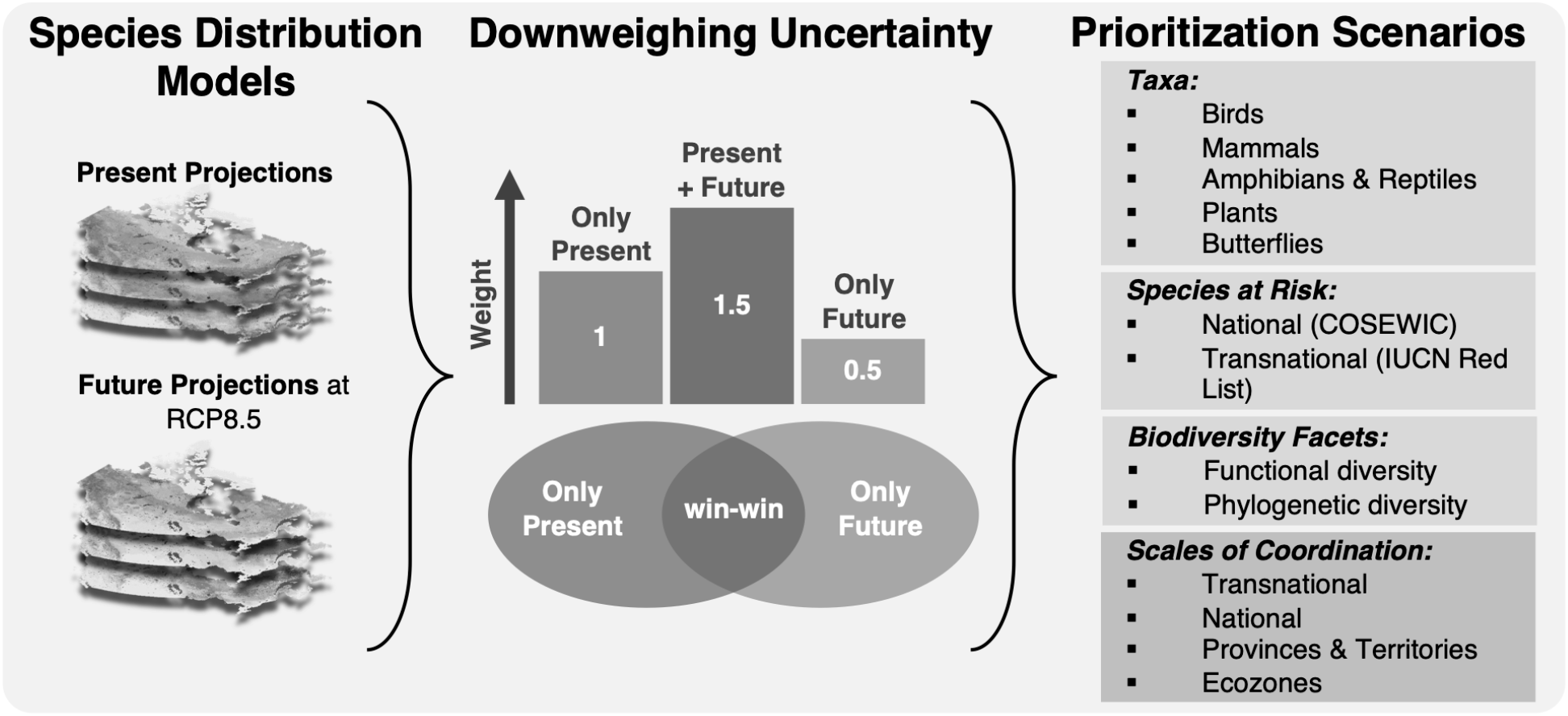
Conceptual methodology. We used current and future projections of species distributions to identify “win-win” areas of climatic stability, which were prioritized across 13 scenarios designed to test how conservation priorities and scales of coordination impact our ability to safeguard species into the future under climate change.

We asked how prioritizing different taxa, species at-risk, and biodiversity facets in spatial planning, as well as how changing the *scale of coordination* (e.g., transnationally, nationally or regionally coordinated) impacts our ability to safeguard biodiversity through 30×30. We designed 13 conservation scenarios, each protecting the same amount of land (30%), but differing based on specific conservation priorities set using Zonation 5 (*22*). To make scenarios realistic, we excluded areas of high human footprint and ceded Indigenous land from our analysis. Our baseline scenario is the *National* scenario, which includes all species weighed equally across kingdoms, and represents the optimal prioritization for Canada. We assessed different conservation priorities using scenarios that prioritize different taxa (birds, mammals, amphibians & reptiles, plants and butterflies), only species at-risk using national (Committee on the Status of Endangered Wildlife in Canada or COSEWIC) and global (IUCN Red List) assessments, or different biodiversity facets (functional and phylogenetic). Finally, we tested how prioritizing land at various spatial scales, representing different *scales of coordination* (transnational, national, or regional by Provinces & Territories or ecozones), influenced spatial priorities and our ability to safeguard biodiversity. To set transnational priorities we weighed species according to the portion of their range in Canada, prioritizing endemics.

For each scenario, we expanded protected areas to reach 30×30 and quantified conservation gains as the percent of total biodiversity protected (weighted endemism) (*23*) and the percent of species considered “safeguarded”, meaning those that met or surpassed their conservation target. Targets were assigned using a modified Species Protection Index (SPI) (*24*), ranging from the rarest species requiring 100% of their Canadian range protected to the most common species requiring only 10%. We calculated biodiversity trade-offs as the percentage difference between the baseline (national scenario) and each other scenario, using both weighted endemism (representing change in percent of protected biodiversity) and SPI (representing change in the percent of safeguarded species). Because weighted endemism and SPI trade-offs were highly correlated (Pearson’s r=0.88, p<0.001), we report SPI although both are made available in the supplementary material (Table S1).

We find that existing protected areas, representing 15.4% of intact terrestrial land, do not effectively capture biodiversity and only safeguard 15.1% of all species, 6.6% of nationally-listed species at-risk, and only 1% of amphibians and reptiles, representing just a single species (*Lithobates sylvaticus*). However, large conservation gains are possible under 30×30 and nationally coordinating protection can safeguard more than 65% of all species, including over 40% of nationally-listed species at-risk and over 60% of amphibians & reptiles (Fig. 2a).

**Fig. 2.**
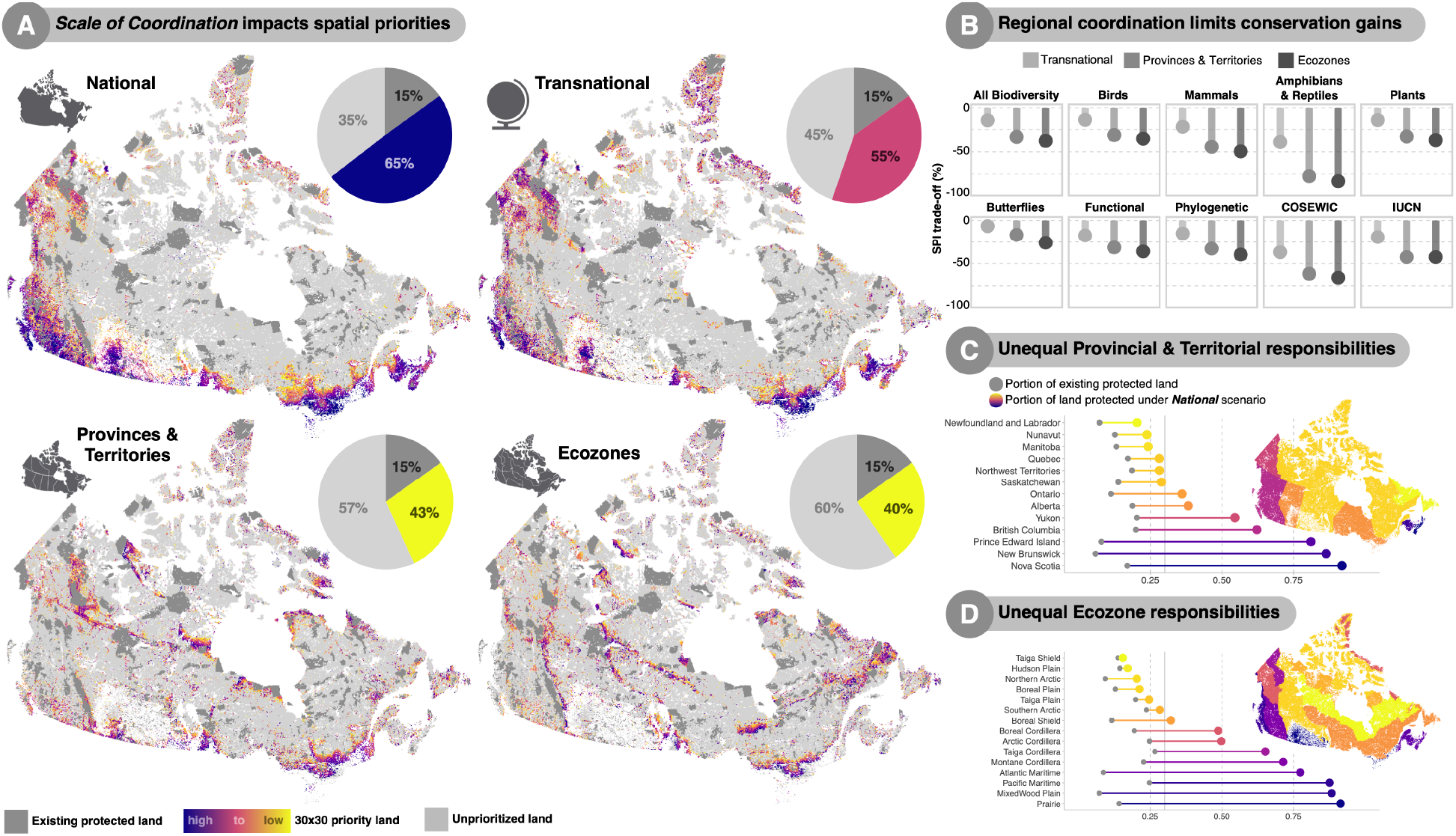
Efficiently protecting biodiversity relies on national coordination. A) 30×30 expansion scenarios across *Scales of Coordination*. Cells are colored by rank, and pies illustrate potential biodiversity gains in the percent of safeguarded species (assessed using SPI), colored by gain. B) Biodiversity SPI trade-offs, representing the percentage loss compared to the baseline *National* scenario, for transnational and regional scenarios across every measure of biodiversity. C-D) The uneven challenge of achieving the baseline *National* scenario captured by the portion of prioritized land across provinces, territories, and ecozones.

Alternative ways of setting priorities incurred trade-offs of varying degrees. Relative to this baseline, scenarios incurred reductions in SPI gains raging from negligible (little to no reduction in the case of *Amphibian & Reptile, Plant, Butterfly, Functional* and *Phylogenetic Diversity*, and *COSEWIC species at-risk* scenarios), to moderate (1-5% reductions in the case of *Mammal*, and *IUCN species at-risk* scenarios), to severe (10-40% reductions in the case of *Bird, Transnational, Provinces & Territories* and *Ecozone* scenarios). Changing the *scale of coordination* by evenly prioritizing land across provinces and territories or ecozones, incurs the largest trade-offs, reducing our ability to safeguard all species by more than 33% and 38% respectively. This decentralized coordination is especially detrimental for safeguarding amphibians, reptiles, and species at-risk (Fig.2b), groups that are already under protected by existing protected land.

While national coordination enables the protection of Canadian biodiversity at large, it relies on highly uneven regional commitments (Fig. 2c-d). Specifically, small maritime provinces, British Columbia, and the arctic Yukon Territory, as well as coastal ecozones and those along Canada’s southern border, contain a high portion of irreplaceable biodiversity. As such, safeguarding Canada’s biodiversity will rely on transboundary coordination and cooperation, demonstrating the uneven challenge of protecting biodiversity through percentage-based targets (*2*, *25*). Interestingly, while regional priorities and spatial representation significantly hinder our ability to safeguard biodiversity, transnational priorities incur smaller trade-offs, suggesting that, at least in Canada, national priorities can efficiently contribute to global goals.

Across all scenarios, spatial priorities varied greatly (Fig. S1), resulting in only a small fraction (~3%) of consistently prioritized land (Fig. 3a). These areas, robust to different conservation priorities, contain over 13% of all biodiversity and over 23% of at-risk biodiversity, making them ideal candidates for protection. A surprisingly high portion of land (~34%) was prioritized in only some scenarios, suggesting much of Canada’s land conservation value is sensitive to shifting conservation priorities (Fig. S2). In general, prioritizing transnational and phylogenetic diversity best reflected the baseline *National* scenario (Fig. 3b), again supporting the idea that prioritizing biodiversity at national scales can serve global goals. *Ecozone, Province & Territory*, and *Amphibian & Reptile* scenarios were least similar to the *National* scenario (Fig. 4a). The spatial discrepancy between taxa-specific scenarios with varying scales of coordination is largely driven by *Amphibian & Reptile* and regional priorities (Fig. 4b), but prioritizing *Amphibians & Reptiles* incurs negligible trade-offs. Overall, regional schemes uncoordinated at the national scale vastly hinders our ability to protect biodiversity at-large (Table S1), indicating that coordinating protection across regions is far more important for the protection of biodiversity at-large than accounting for all species and clades in the spatial planning process.

**Fig. 3.**
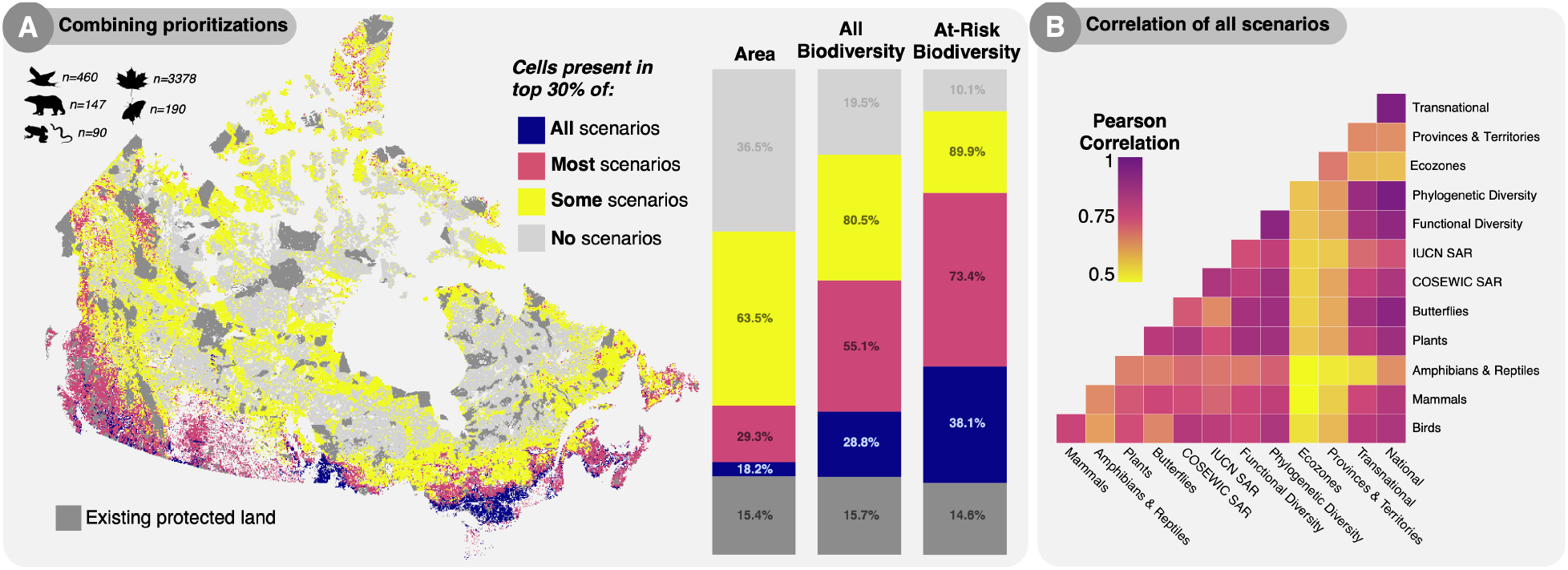
Similarities and differences among priority scenarios. A) Areas important for single or multiple priorities. Bars represent the corresponding percent of land area, biodiversity, and at-risk biodiversity, measured using weighted endemism. B) Correlations of the spatial overlap of across scenarios.

**Fig. 4.**
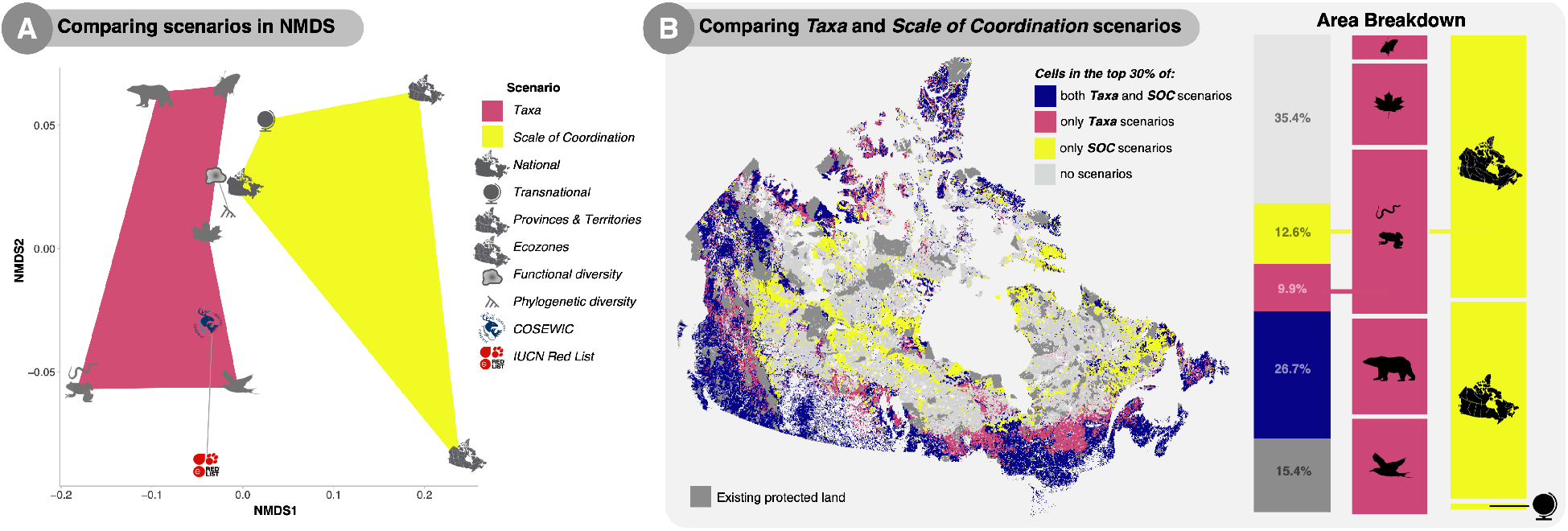
Comparing scenarios reveals amphibian, reptile, and regional priorities drive spatial dissimilarity. A) Comparison of taxa-based and scale of coordination (SOC) scenarios by visualizing the differences in nonmetric multidimensional space (k=2, stress=0.09) and B) spatial overlap. NMDS axes are visualized in Fig. S3.

Protecting Earth’s biodiversity is a global necessity, yet planning predominantly happens at regional and local scales. As such, global biodiversity conservation requires uneven transnational cooperation (*13*), but individual countries have committed to 30×30 and coordinating within a country is far easier than coordinating between countries. Assuming each country meets the goal of 30% protection, our findings show that setting national priorities can efficiently contribute to the protection of global biodiversity (as much as it is possible for an individual country to do so). In Canada, this likely means that transboundary species alone are not responsible for inflated priorities in the south, but that these regions also house a significant portion of endemic taxa. Future work is needed to determine whether similar patterns exist in smaller or more biodiverse countries.

In sum, our results confirm the critical importance of having a biodiversity-informed strategy for 30×30. Scalable biodiversity indicators are essential for understanding the trade-offs associated with different conservation priorities – including which species, functions, and lineages might be left unprotected (*25*). Nationally coordinated strategies will not only most effectively protect the flora and fauna of each country, but can also efficiently contribute to global targets. On the other hand, nationally uncoordinated regional initiatives can limit the ability of area-based conservation to protect biodiversity at large (*26*). Nonetheless, strong arguments can be made for conservation at local scales (*27*), and our findings do not invalidate the potential of such initiatives. Instead, we emphasize the importance of quantifying indicators, assessing trade-offs, and the urgent need for coordinated national strategies for reaching 30×30. The extent to which we can coordinate, cooperate, and measure progress across scales and borders may well determine the success of international targets like 30×30 and consequently, the future of biodiversity on Earth.

## Acknowledgments

The authors would like to acknowledge that while they were able to account for ceded Indigenous land in their analysis, much of what we now call Canada remains unceded, rich with a history of Indigenous oppression. As such, the establishment of new protected areas and Canada’s path to 30×30 should involve Indigenous communities, knowledge, and perspectives, so as to advance Indigenous rights and title. Additionally, the authors would like to thank Darren Li and Cole Lee for their work in gathering the functional trait data, Stefano Mammola for his help with computing functional hypervolumes, and Abbie Gail Jones and Olivia Rahn for their many helpful comments throughout the duration of the project.

## Author contributions

Conceptualization: IE, LJP
Methodology: IE, AB, DC, FR, LJP
Investigation: IE, LJP
Visualization: IE
Funding acquisition: LJP
Project administration: LJP
Supervision: LJP
Writing – original draft: IE
Writing – review & editing: IE, AB, DC, FR, LJP

## Data and materials availability

All data used is publicly available aside from a few range polygons used to constrain species distribution models. Those data, along with our calculated functional, phylogenetic, and transnational weights and all code, as well as scenario rank maps and maps of species richness and endemism are available in the following repository: https://figshare.com/s/0551e56687ba119c7bb8

## Supplementary Materials

Materials and Methods

Figs. S1 to S3

Table S1

