## Supplementary material for "30×30 biodiversity gains rely on national coordination": Methods

### Materials and Methods
#### Species lists and biodiversity data collection

To build species distribution models, we used data for all species downloaded from the Global Biodiversity Informatics Facility (*28*). We first curated species lists for each group. For plants, we used the database of vascular plants of Canada including only native species (*29*). For butterflies, we downloaded from GBIF all observations recorded in Canada with accuracy < 5 km collected between 1990 and 2021, and combined this dataset with all observations available for the same time period in the eButterfly community science platform (www.e-butterfly.com). For terrestrial vertebrates, we first extracted all data from GBIF with a single locality recorded in Canada, then refined the list manually by examining each species removing non-native, extinct and domestic species. We created an index using the 'taxize' in package R (*30*) and manual refinement using a wide range of sources to crosscheck species names between the GBIF backbone taxonomy, IUCN range maps, traits, and phylogenetic data. We then extracted all occurrence points with a geographic location (excluding those with an unidentifiable geographic datum or location details). Due to the starkly different sampling intensities of some groups, we balanced having enough samples with the increased precision of geographic locations in more recent years. This meant that, for birds, we extracted data from 1990 to present and for all other groups, we extracted data from 1970 to present. We further filtered the bird occurrences to capture breeding ranges only (rather than the entire migratory range which often extends well outside of Canada). The occurrence data for birds were further filtered to include the average start and end date of the breeding season (June to July), estimated for a random subset of 25% of species. For species whose data was limited by a lack of associated dates, we included all occurrence points but filtered by IUCN breeding polygons (*31*).

To further clean data points unlikely to represent the native distribution, we removed any points in core urban areas (i.e. areas designated ‘urban or built-up’ in the land-use/land cover data described below), which often included clusters of data points in zoos/sanctuaries. Then, we manually went through each vertebrate species to remove additional points outside of the known range of the species. To do this, we visually compared points to the IUCN range polygons, information about the specific species from guide books or online, and expert knowledge. In some cases, true outlying populations were outside of the range maps, and those populations were retained. For the species flagged as having outlier points, we then used an automated approach for removing outliers for flagged species by removing any points with an average distance to the nearest three other occurrence points of a certain distance in kms (distances established for each group of species independently). We used an average of three rather than one is that some outliers outside the native distribution were recorded multiple times spanning more than one grid cell. Following outlier removal, occurrence data was then gridded for a 1-km$^2$ grid on a Lambert Conformal Conic projection to match the climate data. All data were extracted from GBIF the week of 25-May-2021.

**Climate and edaphic explanatory variables**

We used the following set of climatic variables that were biologically meaningful and had low correlation: mean annual precipitation (mm), chilling degree days (Degree-days below 0°C), precipitation as snow (mm), Hargreave’s climatic moisture index and warming degree-days above 18°C. We used both current and future climate models from Adapt West (*32*). Current climate data is based on PRISM and WorldClim and spans 1991-2020. Future climate projections were downscaled from the Coupled Model Intercomparison Project Phase 6 (CMIP6) based on an ensemble projection from 13 climate models and Representative Concentration

Pathway (RCP 8.5). RCP 8.5 is considered business-as-usual without climate change mitigation, which can produce the most extreme climate change projections. However, most discrepancy in SDM comes from the type of SDM rather than the GCP or RCP except for common species in no dispersal scenarios (*33*). We are also discounting the future and truncating the distance that species can move in future projections (see below), so we felt this evidence justifies the ensemble GCM and RCP 8.5 scenario.

We also used topographic wetness index (calculated based on the 1-km digital elevation model using package 'dynatopmodel' in R (*34*)), topographic ruggedness index (from Adaptwest), and an aggregated land cover layer based on MODIS land cover data and reprojected to our grid and reclassified to: unvegetated, hardwood forests, evergreen forest, mixed forests, shrubs and grasslands (*35*). For plants, we additionally used three variables to represent soil properties (topsoil silt fraction, subsoil pH, and topsoil organic C content) from the Unified North American Soil Map (*36*) (0.25 degree resolution) that were projected to match the 1-km$^2$ climate raster.

**Species distribution models**

We used a set of species distribution models (SDM) that performed well in preliminary tests (variable importance, realistic response curves, visual checks of realistic mapped predictions, and ability to handle interactions between categorical and continuous variables) for a variety of different organisms. All organisms had strongly biased GBIF sampling with the south much more sampled. We addressed this sampling bias by: (1) gridding occurrence data so that a species is either present in the grid cell or not (based on a single occurrence for all non-bird species and 2 occurrences for birds) and (2) accounting for this bias in different ways when fitting models.

We fitted models with three separate algorithms (Generalized Additive Models; GAM, Boosted Regression Trees; BRT, and Maximum Entropy (MaxEnt), which typically have strong predictive power (*37*, *38*). We used two bias correction methods. For BRTs, we fitted models with all environmental predictors plus sample effort (all GBIF observations of plants and vertebrates within a 30-km$^2$ surrounding area) and Human Footprint Index (HFI). Then, we set the sample effort to its maximum value for prediction. MaxEnt uses a target background approach to account for bias (*39*). All presences were used in the model unless they exceeded 5,000, in which case 5000 presences were randomly drawn along with 10,000 absences. While this means there is a presence:absence imbalance for rare species, it was necessary to fully cover the environmental space as we were projecting each species across the study area. Importantly, we are not comparing across species (e.g., calculating richness), Zonation 5 (and the weighted endemism metric more generally) scales all species individually. Therefore, the magnitude of the habitat suitability of one species does not need to be compared to the others. Species with fewer than 10 presences were excluded as were species whose models did not fully converge. Models were bit using the 'dismo' package in R (*40*). Models were fitted on data from all the United States and Canada to avoid environmental truncation and predicted to a 5-km$^2$ grid after initial checks to verify that predicted ranges were very similar between resolutions.

Model validation is a major challenge when true absences are lacking and particularly when we know that input data is strongly biased along the environmental gradients from which we are fitting the model. Our main concern was accounting for this discrepancy, so we validated models based on a set of three comparisons: Canada-wide, colder regions (defined as 80% of the land with the largest Chilling degree days) and warmer regions (the top 20% with the smallest Chilling degree days). We calculated Area under the receiver operator characteristic curve

(AUC) and area under the precision-recall curve (AU-PRC). We did not calculate metrics that require a single threshold as we used the numeric values directly in Zonation. We predicted to both present and future climatic conditions as defined above.

**Mask layers for zonation**

Along with the biodiversity features (projected SDMs), Zonation requires a few other layers. In order to include existing protected areas, and insure those are prioritized first, we used a hierarchical mask layer. We identified existing protected areas using the Canadian Protected and Conserved Areas Database (*21*). Polygons were rasterized and projected to the 1 km$^2$ climate grid, and cells with at least 43% of their area within protected areas were considered protected. We chose to include other effective conservation measures (OECMS) since Canada counts OECMs towards international targets. This threshold was set so that the total number of protected cells would be roughly the same portion as the total amount of protected land (13.5%).

Zonation also accepts a base mask layer that delineates the study area. We wanted to focus our analysis on land in Canada that could feasibly house future protected areas. As such, we excluded areas of high human footprint (representing urban, agricultural, industrial, or other high disturbance areas). We identified areas of high human impact using the recently published Canadian cumulative Human Footprint Index (*19*) reprojected to 1-km$^2$. Following their approach, we considered any cell with an HFI value above 10 to represent an area of “high human footprint”, representing roughly 5.7% of Canada, and excluded it from our analysis. The remaining cells represent largely intact “wilderness” and thus good candidates for protection. Because Canada is largely situated on unceded Indigenous land, we chose to acknowledge what Indigenous land has been ceded by excluding from our analysis, since Indigenous land has been shown to contain levels of biodiversity similar to what is observed in protected areas, and the government has no jurisdiction to establish new protected areas on Indigenous land. To identify Indigenous land, we used the Aboriginal Lands of Canada Legislative Boundaries Database (*41*), rasterized to 1 $km^2$ resolution. We used the same threshold (43%) to identify "Indigenous" cells, representing roughly 6% of Canada.

**Expansion scenarios design**

To assess how different conservation priorities impact our ability to protect biodiversity, we designed 13 conservation scenarios. The national scenario, which represents the optimal scenario for Canada, prioritizes all species simultaneously, weighted equally across kingdoms (so that vertebrate, plant, and butterfly diversity each receive the same total weight). To assess how the inclusion or exclusion of specific taxa impacts spatial priorities, we prioritized birds, mammals, amphibians & reptiles, plants, and butterflies separately. We also designed 2 scenarios to prioritize species-at risk. We used 2 species at risk assessments, the Committee on the Status of Endangered Wildlife in Canada assessment and IUCN's Red List, which correspond to national and global assessments, both accessed in February of 2022. Since, at a national scale, countries are more likely to use their own assessments, we reported the COSEWIC results as the main species at-risk results in the text. For these scenarios, we only included species listed as threatened, endangered, or special concern in the case of COSEWIC and vulnerable, endangered, and critically endangered for IUCN.

To assess conservation scenarios that prioritize functional and phylogenetic biodiversity facets, we weighed species according to their functional or phylogenetic distinctiveness. To calculate functional distinctiveness, we used hypervolume contribution scores. We built a single hypervolume for Canadian vertebrates, plants, and butterflies separately, and calculated the contribution of each species. Contributions were then standardized so the sum of all contributions equaled 1 for vertebrates, plants, and butterflies separately, thus weighing each kingdom evenly during Zonation. To build hypervolumes, we used 2 principal coordinates from a distance matrix calculated from normalized functional traits (*42*) in the 'BAT' package for R (*43*). For vertebrates, we used diet, body mass, litter clutch size, generation length, lifespan, wintering strategy, and age at sexual maturity. Vertebrate functional trait data was sourced from various databases, including the Amniote database (*44*), Amphibio database (*45*), Pantheria database (*46*). For plants, we used seed mass, height, specific leaf area, lifespan, nitrogen fixation capacity, growth form, photosynthetic pathway, dispersal syndrome, reproductive timing, leaf compoundness, and woodiness, all downloaded from the TRY (v5.0) database in February of 2022 (*47*). For butterflies we used expertly estimated mobility, wingspan, range size, and larval host plant breadth from (*48*), combined with the recently compiled LepTraits database, accessed in October of 2022 (*49*). To fill gaps in the trait data, we imputed missing values using phylogenetic vector regressions (PVRs) in the 'PVR' package for R (*50*) calculated from phylogenetic trees to aid random forest imputation in the 'MissForest' package for R (*51*). To calculate phylogenetic distinctiveness, we used existing phylogenetic trees for vertebrates (*52–55*), plants (*56*), and butterflies (*48*). After pruning trees to only include Canadian species, we calculated distinctiveness using the "evol_distinct" command in the 'phyloregion' package for R (*57*).

To assess how spatial scale of coordination impacts spatial priorities, we used three scenarios. The transnational scenario prioritizes global biodiversity by weighing species in Zonation based on their Canadian (weighted) endemism (i.e. the portion of their North-American range found in Canada). The Provinces & Territories scenario protects an even portion of each province and territory, so protected areas are spread evenly across the political landscape. This approach is most similar to what Canada already practices. The Ecozone scenario protects an even portion of each ecozone, achieving even spatial representation from an ecological point of view. Both Province & Territory and Ecozone scenarios represent regional scale priorities.

**Spatial prioritization**

To prioritize land in each conservation scenario, we used Zonation 5 with CAZ2 marginal loss, which balances priorities across species and is new in Zonation 5 (*22*). We chose to exclude areas of high human footprint and Indigenous land from our analysis, as these represent areas where the establishment of new protected land is unlikely (i.e. due to high costs or low availability) or not appropriate. As such, although Canada has protected 13.5% of its terrestrial land, that represents 15.4% of land included in our analysis. We used a hierarchical mask layer in Zonation which allows for the initial prioritization of existing protected land, before prioritizing remaining cells, allowing for complementarity in spatial planning. For Province & Territory and Ecozone runs, we used subregions, where a full prioritization was performed for each subregion separately, and the prioritizations were stitched together in one final raster, representing all of Canada. This allowed each subregion to prioritize the species and endemism specific to that subregion alone, accounting for already protected land, and enabling complementarity. The output of Zonation runs is a final raster for all unmasked cells, ranked according to their priority.

**Reaching 30x30 targets**

From these rank maps, we simulated 30x30 by selecting the top 30% of cells in each scenario, including already protected areas. While all rank maps are available in the supplementary material, we chose to only include the national prioritization as well as the scale of coordination scenarios in the main text.

**Biodiversity trade-offs**

To assess the biodiversity trade-offs associated with different conservation priorities, we used a modified species protection index (SPI) to measure the percentage of taxa considered adequately protected (*24*). SPI works by setting species-specific protection targets, based on how common or rare a species is across the landscape. Since we used probabilistic species distributions, we consider a species range to be the sum of probabilities in all cells across a landscape. The top 10% most common species, with large ranges, require at least 10% of their range inside protected areas to be considered adequately protected. The top 10% rarest species, with small ranges, require 100% of their range inside protected areas to be considered adequately protected. For the 80% of species between these thresholds, we used a log-linear model to set species-specific goals. Species that met or surpassed their conservation goals were considered "protected", while species that did not meet their goals were left "unprotected" by conservation expansion scenarios.

We calculated biodiversity trade-offs as the relative difference between the percentage of species considered "protected" under the *national* prioritization and the number of species considered "protected" under other scenarios. For example, if the national prioritization protected 10 species, and a different conservation priority scenario only protected 5 species, the biodiversity trade-off would be (5/10)*100 = 50%. Or in other words, the different conservation priority protects 50% of the biodiversity compared to the optimum national prioritization. We calculated biodiversity trade-offs for all biodiversity (all species) as well as for birds, mammals, amphibians & reptiles, plants, butterflies, COSEWIC species at-risk, and IUCN species at-risk separately. In addition, we also quantified functional and phylogenetic biodiversity trade-offs by calculating the total functional and phylogenetic contributions of "protected" species as a percentage of total Canadian functional and phylogenetic diversity (the sum of all species contributions).

**Spatial commitments**

To highlight the uneven challenge posed when prioritizing biodiversity, we calculated the total amount of each ecozone and province & territory in the top 30% of the national prioritization scenario.

**Comparing priority scenarios**

To compare scenarios, we used multiple methods. First, we visualized overlap between 30x30 expansion scenarios by highlighting cells in the top 30% of all scenarios, most scenarios (i.e. 7 or more), some scenarios (i.e. 6 or fewer), and no scenarios. Then we calculated the pairwise Pearson correlations for all scenarios and visualized them using a heatmap. Strong correlations represent spatially similar prioritization scenarios, whereas weaker correlations represent spatially divergent scenarios. To compare scenarios further, we use nonmetric multidimensional scaling in the 'vegan' package for R (*58*), treating each prioritization as a site, and each cell as a species. NMDS revealed 2 distinct spatial axes of variation between all scenarios, achieving a low stress value of 0.09. Finally, to highlight the spatial differences between clade and political scenarios, we once again visualized overlap, this time quantifying which specific scenarios were driving the spatial dissimilarity between the two groups
