## Supplementary Figures for "30×30 biodiversity gains rely on national coordination"

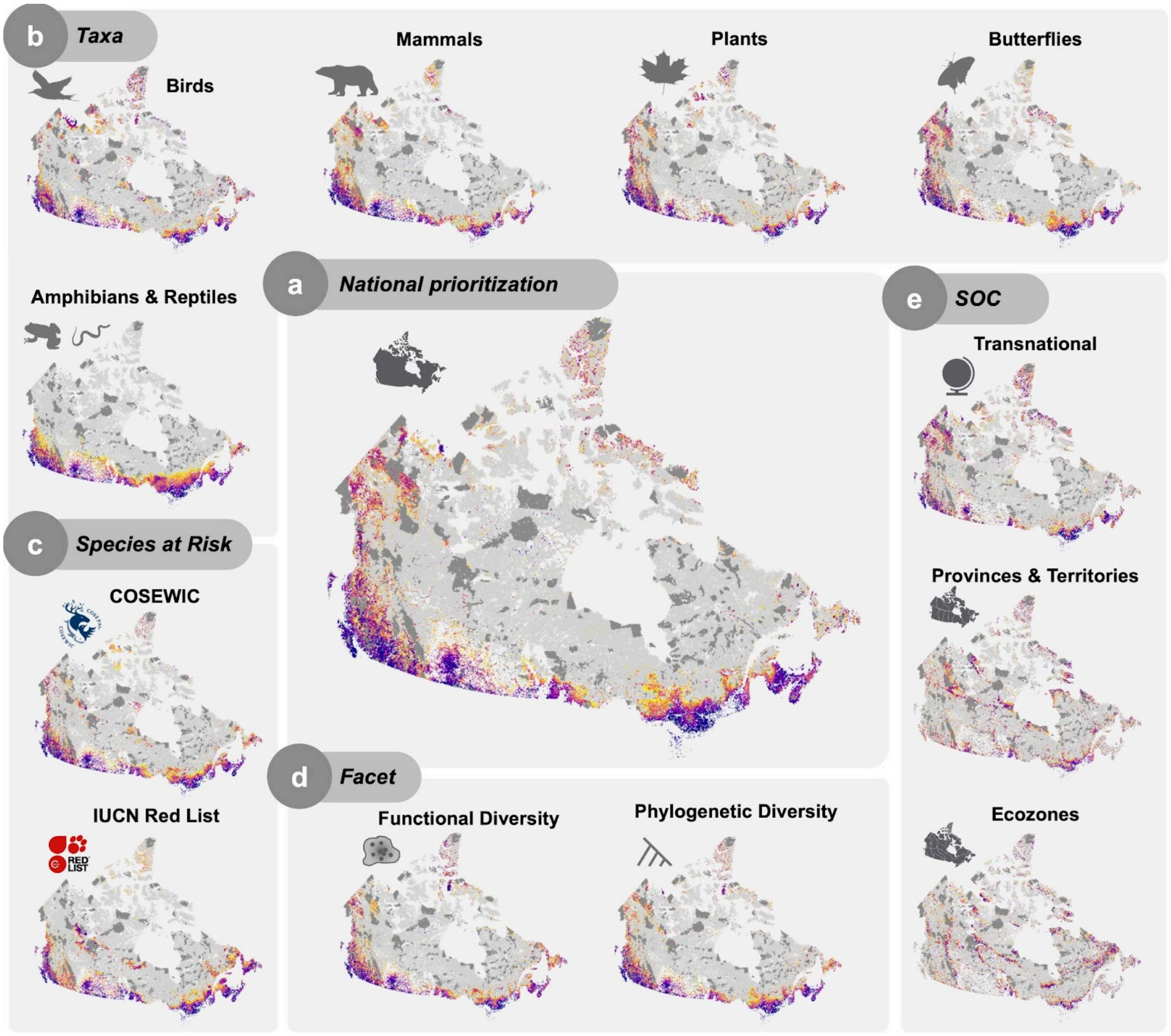


Fig. S1. 30x30 prioritizations for all conservation priorities. The national prioritization (a) represents the optimum scenario for Canada, where all species, weighed equally across kingdoms, are prioritized at the national level, maximizing coverage across taxa. Taxa (b) prioritization scenarios include specific clades, species at-risk (c) prioritization scenarios include only species listed as at-risk by COSEWIC or IUCN Red List, and Facet (d) prioritization scenarios prioritize functional and phylogenetic diversity. Scale of Coordination (SOC) (e) prioritization scenarios prioritize land differently across spatial scales, either prioritizing transnational diversity using species-specific weights based on endemism or provincial &

territorial and ecozone diversity by performing separate prioritizations for each region, thereby achieving spatial representation. Land in the top 30% of each prioritization scenario is highlighted in color with blue cells representing the most important land, followed by red, and then yellow. Existing protected areas are colored in dark grey, and light grey land represents the 70% of Canada that falls outside 30x30 spatial priorities.

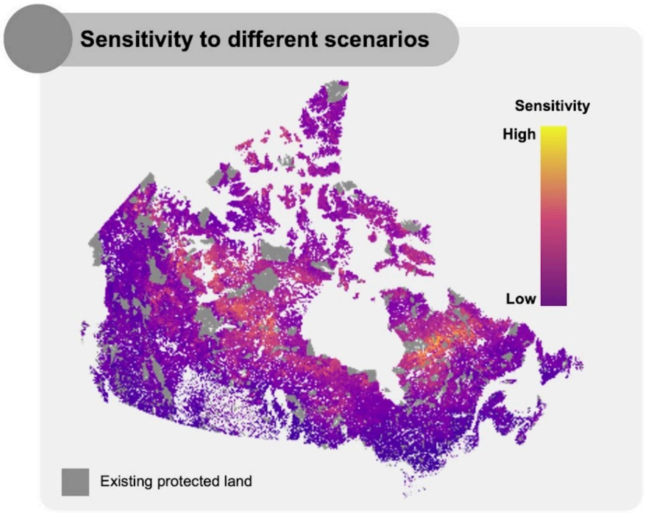


**Fig. S2.** Spatial sensitivity of land to shifting conservation priorities. We calculated sensitivity as the standard deviation of cell rank across all prioritization scenarios divided by the mean. Existing protected areas are colored in dark grey. In general, sensitivity was lower in southern Canada, along the coasts, and in the arctic. Sensitivity was high in the interior boreal region, due to a combination of lower cell ranks in general and sensitivity to specific conservation priorities such as Provinces & Territories and Ecozone priorities.

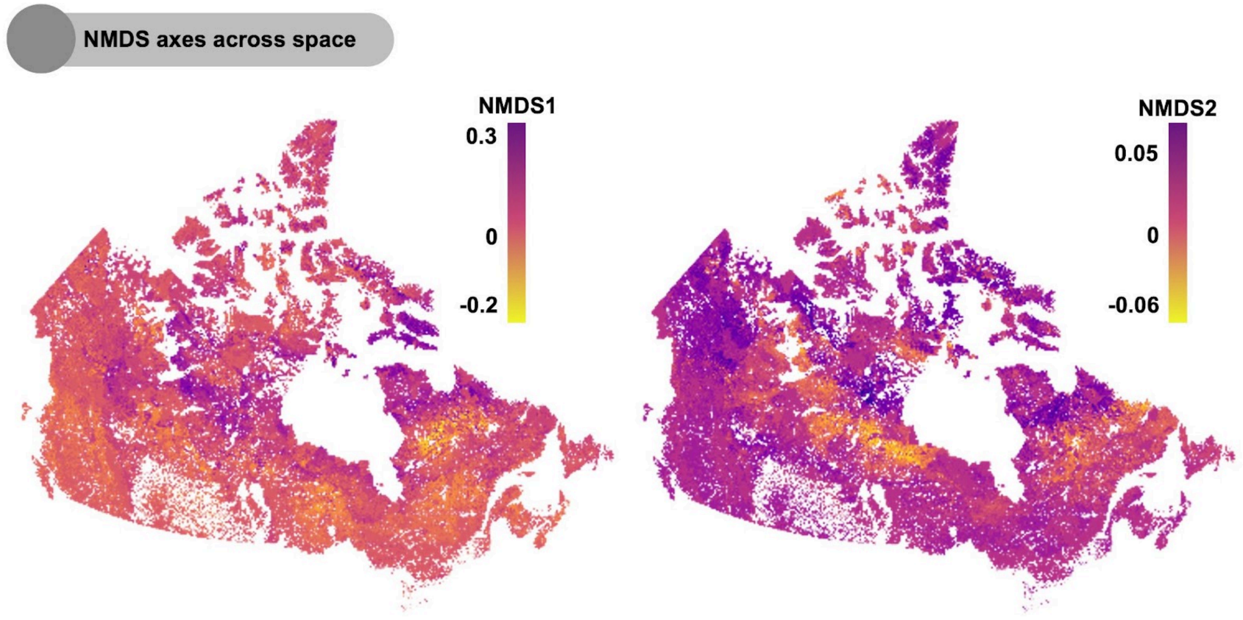


**Fig. S3.** NMDS axes 1 and 2 mapped across space. These axes correspond to those in Fig. 4 and represent the top axes of variation in cell rank across all conservation prioritizations.
