## Supplementary Tables for "30×30 biodiversity gains rely on national coordination"

| Priority | Priority Group | Biodiversity Measure | Already Protected | 30x30 Gain | Trade-off | Already Protected | 30x30 Gain | Trade-off |
|---|---|---|---|---|---|---|---|---|
| | | | Species Protection Index (%) | | | Weighted Endemism (%) | | |
| **National** | **Scale of Coordination** | **Total Biodiversity** | **15.1** | **64.5** | **-** | **15.7** | **59.2** | **-** |
| National | Scale of Coordination | Birds | 20.6 | 76.9 | - | 15.5 | 52.9 | - |
| National | Scale of Coordination | Mammals | 14.3 | 65.9 | - | 16.7 | 61.4 | - |
| National | Scale of Coordination | Amphibians & Reptiles | 1.1 | 60.7 | - | 13.1 | 78.2 | - |
| National | Scale of Coordination | Plants | 14.8 | 63.6 | - | 16.3 | 64.7 | - |
| National | Scale of Coordination | Butterflies | 14.3 | 54.8 | - | 15.3 | 54.8 | - |
| National | Scale of Coordination | COSEWIC SAR | 6.6 | 40.5 | - | 14.6 | 75.4 | - |
| National | Scale of Coordination | IUCN SAR | 13.9 | 60.5 | - | 14.6 | 61.8 | - |
| National | Scale of Coordination | Functional Diversity | 19.6 | 70.2 | - | 15.2 | 59.3 | - |
| National | Scale of Coordination | Phylogenetic Diversity | 15.0 | 64.7 | - | 15.4 | 60.7 | - |
| **Transnational** | **Scale of Coordination** | **Total Biodiversity** | **15.1** | **55.3** | **-14.3** | **15.7** | **55.6** | **-6.1** |
| Transnational | Scale of Coordination | Birds | 20.6 | 66.4 | -13.7 | 15.5 | 49.1 | -7.2 |
| Transnational | Scale of Coordination | Mammals | 14.3 | 51.7 | -21.5 | 16.7 | 56.2 | -8.5 |
| Transnational | Scale of Coordination | Amphibians & Reptiles | 1.1 | 37.1 | -38.9 | 13.1 | 71.1 | -9.1 |
| Transnational | Scale of Coordination | Plants | 14.8 | 54.8 | -13.8 | 16.3 | 61.1 | -5.6 |
| Transnational | Scale of Coordination | Butterflies | 14.3 | 51.2 | -6.6 | 15.3 | 52.2 | -4.7 |
| Transnational | Scale of Coordination | COSEWIC SAR | 6.6 | 25.6 | -36.8 | 14.6 | 70.2 | -6.9 |
| Transnational | Scale of Coordination | IUCN SAR | 13.9 | 48.8 | -19.3 | 14.6 | 58.0 | -6.1 |
| Transnational | Scale of Coordination | Functional Diversity | 19.6 | 57.9 | -17.5 | 15.2 | 55.4 | -6.6 |
| Transnational | Scale of Coordination | Phylogenetic Diversity | 15.0 | 54.9 | -15.1 | 15.4 | 56.7 | -6.6 |
| **Provinces & Territories** | **Scale of Coordination** | **Total Biodiversity** | **15.1** | **43.1** | **-33.2** | **15.7** | **47.2** | **-20.3** |
| Provinces & Territories | Scale of Coordination | Birds | 20.6 | 53.0 | -31.1 | 15.5 | 43.2 | -18.3 |
| Provinces & Territories | Scale of Coordination | Mammals | 14.3 | 36.7 | -44.3 | 16.7 | 46.6 | -24.1 |
| Provinces & Territories | Scale of Coordination | Amphibians & Reptiles | 1.1 | 13.5 | -77.8 | 13.1 | 59.6 | -23.8 |
| Provinces & Territories | Scale of Coordination | Plants | 14.8 | 42.8 | -32.7 | 16.3 | 51.1 | -21.0 |
| Provinces & Territories | Scale of Coordination | Butterflies | 14.3 | 45.6 | -16.8 | 15.3 | 44.4 | -19.0 |
| Provinces & Territories | Scale of Coordination | COSEWIC SAR | 6.6 | 15.6 | -61.5 | 14.6 | 59.5 | -21.1 |
| Provinces & Territories | Scale of Coordination | IUCN SAR | 13.9 | 34.9 | -42.3 | 14.6 | 50.4 | -18.4 |
| Provinces & Territories | Scale of Coordination | Functional Diversity | 19.6 | 48.2 | -31.3 | 15.2 | 47.8 | -19.4 |
| Provinces & Territories | Scale of Coordination | Phylogenetic Diversity | 15.0 | 43.5 | -32.8 | 15.4 | 48.0 | -20.9 |
| **Ecozones** | **Scale of Coordination** | **Total Biodiversity** | **15.1** | **40.3** | **-37.5** | **15.7** | **41.7** | **-29.6** |
| Ecozones | Scale of Coordination | Birds | 20.6 | 49.9 | -35.1 | 15.5 | 38.6 | -27.0 |

| | | | | | | | | |
|---|---|---|---|---|---|---|---|---|
| Ecozones | Scale of Coordination | Mammals | 14.3 | 33.3 | -49.5 | 16.7 | 39.3 | -36.0 |
| Ecozones | Scale of Coordination | Amphibians & Reptiles | 1.1 | 10.1 | -83.4 | 13.1 | 47.3 | -39.5 |
| Ecozones | Scale of Coordination | Plants | 14.8 | 40.2 | -36.8 | 16.3 | 44.3 | -31.5 |
| Ecozones | Scale of Coordination | Butterflies | 14.3 | 40.6 | -25.9 | 15.3 | 40.9 | -25.4 |
| Ecozones | Scale of Coordination | COSEWIC SAR | 6.6 | 13.8 | -65.9 | 14.6 | 47.0 | -37.7 |
| Ecozones | Scale of Coordination | IUCN SAR | 13.9 | 34.9 | -42.3 | 14.6 | 44.3 | -28.3 |
| Ecozones | Scale of Coordination | Functional Diversity | 19.6 | 44.9 | -36 | 15.2 | 41.7 | -29.7 |
| Ecozones | Scale of Coordination | Phylogenetic Diversity | 15.0 | 38.9 | -39.9 | 15.4 | 41.9 | -31.0 |
| **Birds** | **Taxa** | **Total Biodiversity** | **15.1** | **57.5** | **-10.9** | **15.7** | **55.8** | **-5.7** |
| Birds | Taxa | Birds | 20.6 | 77.4 | 0.7 | 15.5 | 53.2 | 0.6 |
| Birds | Taxa | Mammals | 14.3 | 57.8 | -12.3 | 16.7 | 58.4 | -4.9 |
| Birds | Taxa | Amphibians & Reptiles | 1.1 | 55.1 | -9.2 | 13.1 | 75.6 | -3.3 |
| Birds | Taxa | Plants | 14.8 | 55.8 | -12.3 | 16.3 | 60.7 | -6.2 |
| Birds | Taxa | Butterflies | 14.3 | 43.8 | -20.1 | 15.3 | 49.5 | -9.7 |
| Birds | Taxa | COSEWIC SAR | 6.6 | 35.3 | -12.8 | 14.6 | 73.2 | -2.9 |
| Birds | Taxa | IUCN SAR | 13.9 | 58.1 | -4 | 14.6 | 61.6 | -0.3 |
| Birds | Taxa | Functional Diversity | 19.6 | 64.1 | -8.7 | 15.2 | 56.3 | -5.1 |
| Birds | Taxa | Phylogenetic Diversity | 15.0 | 56.8 | -12.2 | 15.4 | 57.6 | -5.1 |
| **Mammals** | **Taxa** | **Total Biodiversity** | **15.1** | **63.7** | **-1.2** | **15.7** | **58.6** | **-1.0** |
| Mammals | Taxa | Birds | 20.6 | 76.9 | 0 | 15.5 | 53.3 | 0.8 |
| Mammals | Taxa | Mammals | 14.3 | 66.7 | 1.2 | 16.7 | 63.9 | 4.1 |
| Mammals | Taxa | Amphibians & Reptiles | 1.1 | 62.9 | 3.6 | 13.1 | 78.3 | 0.1 |
| Mammals | Taxa | Plants | 14.8 | 62.6 | -1.6 | 16.3 | 63.9 | -1.2 |
| Mammals | Taxa | Butterflies | 14.3 | 52.1 | -4.9 | 15.3 | 53.1 | -3.1 |
| Mammals | Taxa | COSEWIC SAR | 6.6 | 39.4 | -2.7 | 14.6 | 74.9 | -0.7 |
| Mammals | Taxa | IUCN SAR | 13.9 | 58.1 | -4 | 14.6 | 60.8 | -1.6 |
| Mammals | Taxa | Functional Diversity | 19.6 | 68.9 | -1.9 | 15.2 | 58.8 | -0.8 |
| Mammals | Taxa | Phylogenetic Diversity | 15.0 | 63.2 | -2.3 | 15.4 | 60.1 | -1.0 |
| **Amphibians & Reptiles** | **Taxa** | **Total Biodiversity** | **15.1** | **69.9** | **8.4** | **15.7** | **60.2** | **1.7** |
| Amphibians & Reptiles | Taxa | Birds | 20.6 | 78.9 | 2.6 | 15.5 | 55.8 | 5.5 |
| Amphibians & Reptiles | Taxa | Mammals | 14.3 | 68.7 | 4.2 | 16.7 | 63.0 | 2.6 |
| Amphibians & Reptiles | Taxa | Amphibians & Reptiles | 1.1 | 79.8 | 31.5 | 13.1 | 88.1 | 12.7 |
| Amphibians & Reptiles | Taxa | Plants | 14.8 | 69.4 | 9.1 | 16.3 | 65.3 | 0.9 |
| Amphibians & Reptiles | Taxa | Butterflies | 14.3 | 55.3 | 0.9 | 15.3 | 53.9 | -1.6 |
| Amphibians & Reptiles | Taxa | COSEWIC SAR | 6.6 | 59.5 | 46.9 | 14.6 | 79.9 | 6 |
| Amphibians & Reptiles | Taxa | IUCN SAR | 13.9 | 67.4 | 11.4 | 14.6 | 66.7 | 7.9 |

| | | | | | | | | |
|---|---|---|---|---|---|---|---|---|
| Amphibians & Reptiles | Taxa | Functional Diversity | 19.6 | 74.7 | 6.4 | 15.2 | 61.4 | 3.5 |
| Amphibians & Reptiles | Taxa | Phylogenetic Diversity | 15.0 | 68.5 | 5.9 | 15.4 | 62.9 | 3.6 |
| **Plants** | **Taxa** | **Total Biodiversity** | **15.1** | **64.8** | **0.5** | **15.7** | **58.9** | **-0.5** |
| Plants | Taxa | Birds | 20.6 | 73.6 | -4.3 | 15.5 | 52.2 | -1.3 |
| Plants | Taxa | Mammals | 14.3 | 62.5 | -5.2 | 16.7 | 60.5 | -1.5 |
| Plants | Taxa | Amphibians & Reptiles | 1.1 | 53.9 | -11.2 | 13.1 | 76.6 | -2.0 |
| Plants | Taxa | Plants | 14.8 | 64.9 | 2 | 16.3 | 66.2 | 2.3 |
| Plants | Taxa | Butterflies | 14.3 | 51.2 | -6.6 | 15.3 | 53.5 | -2.4 |
| Plants | Taxa | COSEWIC SAR | 6.6 | 36.3 | -10.4 | 14.6 | 75.1 | -0.4 |
| Plants | Taxa | IUCN SAR | 13.9 | 55.8 | -7.8 | 14.6 | 62.8 | 1.6 |
| Plants | Taxa | Functional Diversity | 19.6 | 67.8 | -3.4 | 15.2 | 58.6 | -1.2 |
| Plants | Taxa | Phylogenetic Diversity | 15.0 | 62.3 | -3.7 | 15.4 | 60.5 | -0.3 |
| **Butterflies** | **Taxa** | **Total Biodiversity** | **15.1** | **64.6** | **0.2** | **15.7** | **59.2** | **0.0** |
| Butterflies | Taxa | Birds | 20.6 | 74.5 | -3.1 | 15.5 | 51.9 | -1.9 |
| Butterflies | Taxa | Mammals | 14.3 | 63.9 | -3 | 16.7 | 60.5 | -1.5 |
| Butterflies | Taxa | Amphibians & Reptiles | 1.1 | 60.7 | 0 | 13.1 | 77.7 | -0.6 |
| Butterflies | Taxa | Plants | 14.8 | 64.0 | 0.6 | 16.3 | 64.6 | -0.2 |
| Butterflies | Taxa | Butterflies | 14.3 | 56.2 | 2.6 | 15.3 | 55.9 | 2.0 |
| Butterflies | Taxa | COSEWIC SAR | 6.6 | 40.1 | -1 | 14.6 | 74.7 | -0.9 |
| Butterflies | Taxa | IUCN SAR | 13.9 | 53.5 | -11.6 | 14.6 | 59.9 | -3.1 |
| Butterflies | Taxa | Functional Diversity | 19.6 | 70.9 | 1 | 15.2 | 59.1 | -0.3 |
| Butterflies | Taxa | Phylogenetic Diversity | 15.0 | 64.7 | 0 | 15.4 | 60.6 | -0.2 |
| **COSEWIC SAR** | **Taxa** | **Total Biodiversity** | **15.1** | **66.1** | **2.5** | **15.7** | **58.6** | **-1.0** |
| COSEWIC SAR | Taxa | Birds | 20.6 | 78.9 | 2.6 | 15.5 | 53.4 | 0.9 |
| COSEWIC SAR | Taxa | Mammals | 14.3 | 65.3 | -0.9 | 16.7 | 60.8 | -1.0 |
| COSEWIC SAR | Taxa | Amphibians & Reptiles | 1.1 | 68.5 | 12.9 | 13.1 | 81.4 | 4.1 |
| COSEWIC SAR | Taxa | Plants | 14.8 | 65.4 | 2.8 | 16.3 | 64.4 | -0.5 |
| COSEWIC SAR | Taxa | Butterflies | 14.3 | 49.3 | -10 | 15.3 | 64.5 | 17.7 |
| COSEWIC SAR | Taxa | COSEWIC SAR | 6.6 | 46.7 | 15.3 | 14.6 | 77.7 | 3.1 |
| COSEWIC SAR | Taxa | IUCN SAR | 13.9 | 58.1 | -4 | 14.6 | 64.5 | 4.4 |
| COSEWIC SAR | Taxa | Functional Diversity | 19.6 | 69.4 | -1.1 | 15.2 | 59.0 | -0.5 |
| COSEWIC SAR | Taxa | Phylogenetic Diversity | 15.0 | 63.9 | -1.2 | 15.4 | 60.6 | -0.2 |
| **IUCN SAR** | **Taxa** | **Total Biodiversity** | **15.1** | **61.7** | **-4.3** | **15.7** | **56.2** | **-5.1** |
| IUCN SAR | Taxa | Birds | 20.6 | 76.9 | 0 | 15.5 | 52.1 | -1.5 |
| IUCN SAR | Taxa | Mammals | 14.3 | 58.5 | -11.2 | 16.7 | 57.9 | -5.7 |
| IUCN SAR | Taxa | Amphibians & Reptiles | 1.1 | 55.1 | -9.2 | 13.1 | 78.3 | 0.1 |

| | | | | | | | | |
|---|---|---|---|---|---|---|---|---|
| IUCN SAR | Taxa | Plants | 14.8 | 61.2 | -3.8 | 16.3 | 62.3 | -3.7 |
| IUCN SAR | Taxa | Butterflies | 14.3 | 45.2 | -17.5 | 15.3 | 49.5 | -9.7 |
| IUCN SAR | Taxa | COSEWIC SAR | 6.6 | 38.4 | -5.2 | 14.6 | 74.6 | -1.1 |
| IUCN SAR | Taxa | IUCN SAR | 13.9 | 60.5 | 0 | 14.6 | 67.2 | 8.7 |
| IUCN SAR | Taxa | Functional Diversity | 19.6 | 65.6 | -6.6 | 15.2 | 56.9 | -4.0 |
| IUCN SAR | Taxa | Phylogenetic Diversity | 15.0 | 60.1 | -7.1 | 15.4 | 58.5 | -3.6 |
| **Functional Diversity** | **Facet** | **Total Biodiversity** | **15.1** | **65.5** | **1.6** | **15.7** | **58.9** | **-0.5** |
| Functional Diversity | Facet | Birds | 20.6 | 76.5 | -0.5 | 15.5 | 53.0 | 0.2 |
| Functional Diversity | Facet | Mammals | 14.3 | 65.3 | -0.9 | 16.7 | 61.3 | -0.2 |
| Functional Diversity | Facet | Amphibians & Reptiles | 1.1 | 61.8 | 1.8 | 13.1 | 78.8 | 0.8 |
| Functional Diversity | Facet | Plants | 14.8 | 64.9 | 2 | 16.3 | 64.7 | 0 |
| Functional Diversity | Facet | Butterflies | 14.3 | 53.5 | -2.4 | 15.3 | 53.8 | -1.8 |
| Functional Diversity | Facet | COSEWIC SAR | 6.6 | 42.9 | 5.9 | 14.6 | 75.8 | 0.5 |
| Functional Diversity | Facet | IUCN SAR | 13.9 | 60.5 | 0 | 14.6 | 63.1 | 2.1 |
| Functional Diversity | Facet | Functional Diversity | 19.6 | 70.3 | 0.1 | 15.2 | 59.6 | 0.5 |
| Functional Diversity | Facet | Phylogenetic Diversity | 15.0 | 64.9 | 0.3 | 15.4 | 60.7 | 0.0 |
| **Phylogenetic Diversity** | **Facet** | **Total Biodiversity** | **15.1** | **67.9** | **5.3** | **15.7** | **59.7** | **0.8** |
| Phylogenetic Diversity | Facet | Birds | 20.6 | 78.5 | 2.1 | 15.5 | 53.8 | 1.7 |
| Phylogenetic Diversity | Facet | Mammals | 14.3 | 69.4 | 5.3 | 16.7 | 61.6 | 0.3 |
| Phylogenetic Diversity | Facet | Amphibians & Reptiles | 1.1 | 68.5 | 12.9 | 13.1 | 80.8 | 3.3 |
| Phylogenetic Diversity | Facet | Plants | 14.8 | 67.3 | 5.8 | 16.3 | 65.3 | 0.9 |
| Phylogenetic Diversity | Facet | Butterflies | 14.3 | 53.9 | -1.6 | 15.3 | 54.7 | -0.2 |
| Phylogenetic Diversity | Facet | COSEWIC SAR | 6.6 | 48.4 | 19.5 | 14.6 | 76.9 | 2 |
| Phylogenetic Diversity | Facet | IUCN SAR | 13.9 | 58.1 | -4 | 14.6 | 64.1 | 3.7 |
| Phylogenetic Diversity | Facet | Functional Diversity | 19.6 | 71.9 | 2.4 | 15.2 | 60.0 | 1.2 |
| Phylogenetic Diversity | Facet | Phylogenetic Diversity | 15.0 | 66.3 | 2.5 | 15.4 | 61.6 | 1.5 |

**Table S1.** Different conservation priorities protect different amounts of biodiversity. Table includes all conservation priorities, and reports the conservation gains possible under 30x30 as the percentage of that measure of biodiversity that would be adequately protected. Trade-offs were calculated as the difference between the potential for each conservation priority and the maximum potential for the national prioritization. A negative trade-off means that that conservation priority limits our ability to protect biodiversity compared to the national prioritization, while a positive trade-off indicates that that priority may protect biodiversity either as well, or better than the national prioritization. Positive trade-offs are likely the result of Zonation's marginal loss rule (CAZ2) that balances spatial priorities across species according to rarity and endemism. So, while we might be able to adequately protect more species by prioritizing amphibian & reptiles in spatial planning, the conservation gains would not be balanced across all taxa (for instance more northern species would be very poorly protected). Weighted endemism and SPI trade-offs were highly correlated (Pearson's $R=0.88$, $p<0.001$). Bolded rows indicate the trade-offs for biodiversity assessed at the national scale for all species and are those references in the main text.
